# BioWardrobe: an integrated platform for analysis of epigenomics and transcriptomics data

**DOI:** 10.1101/012799

**Authors:** Andrey V. Kartashov, Artem Barski

## Abstract

BioWardrobe is an integrated platform that empowers users to store, visualize and analyze epigenomics and transcriptomics data, without the need for programming expertise, by using a biologist-friendly webbased user interface. Predefined pipelines automate download of data from core facilities or public databases, calculate RPKMs and identify peaks and display results via the interactive web interface. Additional capabilities include analyzing differential gene expression and DNA-protein binding and creating average tag density profiles and heatmaps.

---

The recent proliferation of next-generation sequencing (NGS) –based methods for analysis of gene expression, chromatin structure and protein-DNA interactions has opened new horizons for molecular biology. These methods include RNA sequencing (RNA-Seq)^1^, chromatin immunoprecipitation sequencing (ChIP-Seq)^2^, DNase I sequencing (DNase-Seq)^3^, micrococcal nuclease sequencing (MNase-Seq)^4^, assay for transposase-accessible chromatin sequencing (ATAC-Seq)^5^, and others. On the “wet lab” side, these methods are largely well established and can be performed by experienced molecular biologists; however, analysis of the sequenced data requires bioinformatics expertise that many molecular biologists do not possess. Re-utilizing published datasets is also challenging: although authors usually comply with the longstanding requirement to deposit raw data files into databases such as Sequence Read Archive (SRA) or Gene Expression Omnibus (GEO), it is impossible to analyze these datasets without special expertise. Even when processed data files (e.g., gene expression values) are available, direct comparison between datasets is ill advised because different laboratories use different pipelines (or different software versions). This situation means that biologists require the help of bioinformaticians even for the simplest of tasks, such as viewing their own data on a genome browser, putting these exciting techniques beyond the reach of many laboratories. Even when bioinformaticians are available, differences in priorities within collaborations can result in delays and misunderstandings that are damaging to the research effort. An optimal way to mitigate these problems is to enable biologists to perform at least basic tasks without the help of bioinformaticians by providing them with a user friendly data analysis software.

Multiple stand-alone programs and web services are available for the analysis of NGS data. However, the majority of currently available tools have a command-line interface, solve one specific and typically require file conversions between them. The commercial programs Genespring ^6^, Partek ^7^ and Golden Helix ^8^ can be run on regular desktop computers and allow analysis of gene expression or genetic variation. However, users have to load the data manually and store it on their desktop computers; given the sheer volume of NGS datasets, this set up makes data analysis complicated at best. Furthermore, these tools do not allow for seamless integration of multiple, published or locally produced datasets. Illumina Basespace ^9^ and Galaxy server ^10^ allow for both storage and analysis of data and have integrated viewing tools. However, they require transfer of data outside the institution (which may be prohibited by HIPAA regulations in some cases) and provide only limited storage space for user data. Although Galaxy provides the opportunity to run tools without using a command-line interface, users still have to manage file type conversions and select detailed parameters each time, which requires a deep understanding of each tool and file format. Absence of stable pipelines may result in inexperienced users comparing “apples to oranges”. In summary, few of the available tools provide a biologist-friendly interface, and none integrate such an interface with data storage, display and analysis.

We therefore developed BioWardrobe—a biologist-friendly platform for integrated acquisition, storage, display and analysis of NGS data, aimed primarily at researchers in the epigenomics field. BioWardrobe features include download of raw data from core facilities or online databases (e.g., GEO), read mapping and data display on a local instance of the UCSC genome browser ^11^, quality control and both basic and advanced data analysis (**Fig. 1a**). In basic analysis (**Supplementary Fig. 1a**), automated pipelines are used to process each experiment. The pipelines are selected on the basis of biologist-friendly experimental parameters (e.g., RNA/ChIP-Seq, paired/single, genome, stranded/unstranded, antibody) and combine the tools developed by ourselves and by others (e.g., Bowtie ^12^, STAR ^13^, FASTX ^14^ and MACS2 ^15^) with wrappers that enhance the output of original software by offering additional information, provide experimentally meaningful quality controls and display results in a graphical user interface. The quality controls produced during basic analysis were chosen to facilitate troubleshooting of experimental procedures. Customizable advanced analysis can combine multiple experiments and includes tools for comparing gene expression (DESeq1/2 ^16^) and genome occupancy (MAnorm ^17^) profiles between samples or groups of samples and creating principal component analysis plots, gene lists, average tag density profiles and heatmaps using a graphical user interface (**Supplementary Fig. 1b**). Incorporating additional custom scripts is facilitated by a built-in interface for the R programming language. All of the precomputed data are stored in an SQL database and can be accessed via a convenient web interface by biologists. Bioinformaticians, on the other hand, can access the data from R using a provided R library or using other programming languages with standard MySQL frameworks. BioWardrobe can be run on Linux or MacOSX systems (e.g., a Mac Pro desktop). Source code and installation instructions are available at http://biowardrobe.com. A limited-functionality demo version that contains the two dataset discussed below is available at http://demo.biowardrobe.com.

**Figure 1.**
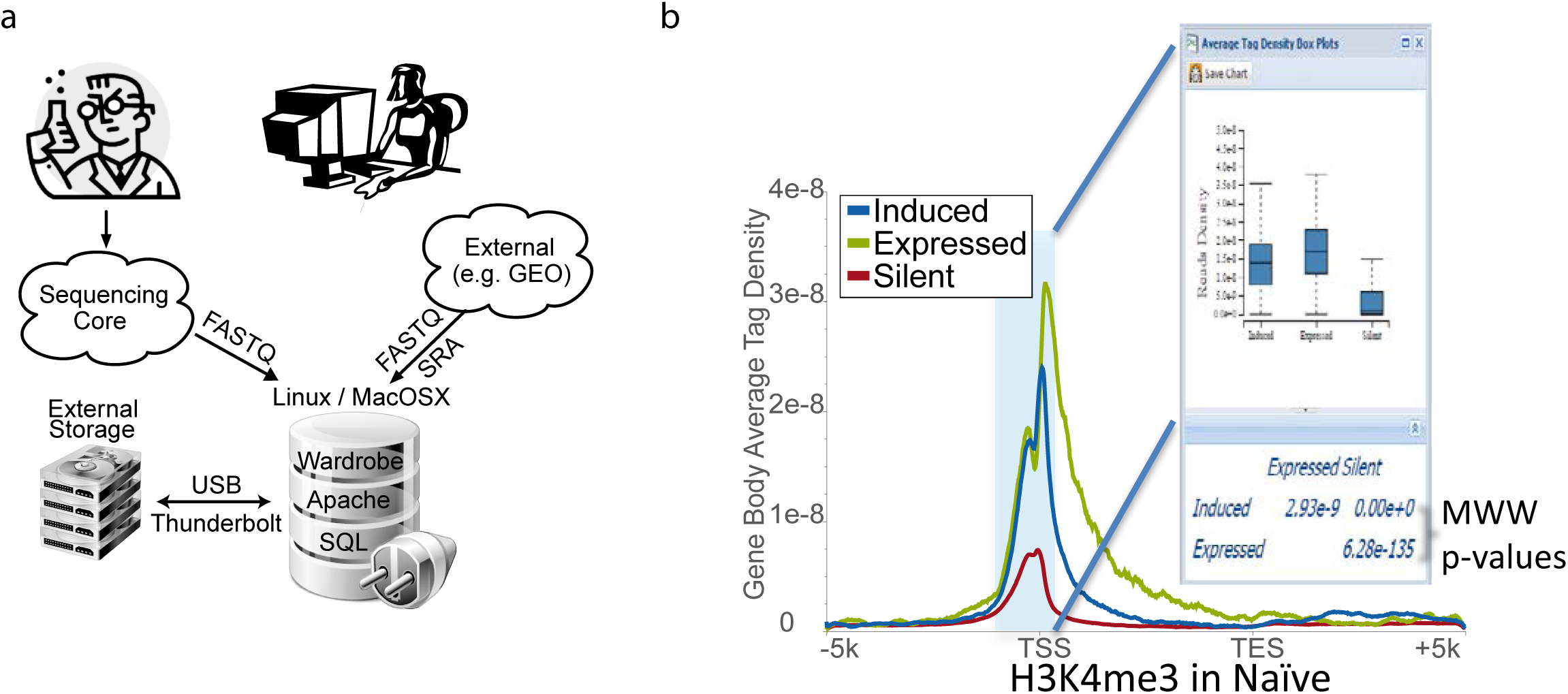
BioWardrobe overview and poising of genes in T cells. **(a)** The BioWardrobe server can be set up on a Linux or Mac computer attached to a consumer-class storage array, typically within a local institutional network. Researchers can use BioWardrobe to upload data from a sequencing core or a public database and promptly receive quality control data, view the results in the browser and perform some of the analysis without the assistance of bioinformaticians. Bioinformaticians can access the precomputed data in Wardrobe’s SQL database to perform further analysis. **(b)** Shown are the H3K4me3 average tag density profiles for the gene body in naïve T cells for genes that are expressed or silent in both Thn and Th1 cells and those that induced during Thn to Th1 transition. The box plot window shows the distribution of H3K4me3 tag densities for the three gene sets within the area shaded in the plot (Silent, Expressed, Induced). MWW p-values are shown below the box plot. Here and in other figures the plot was produced in BioWardrobe, saved as an. svg and adjusted in Illustrator.

To demonstrate the utility of the included quality controls and the ability of BioWardrobe to integrate and analyze data from various sources, we have performed re-analysis of two published datasets. The first study examined gene expression and chromatin changes during differentiation of human naïve T cells (Thn) into T helper type 1 (Th1) cells (SRA082670 ^18^). The dataset included Helicos RNA-Seq performed in triplicates for both resting Thn cells and cells differentiated in Th1 conditions for 72 hours (Th1 cells) and histone 3 lysine residue 4 trimethlation) (H3K4me3) ChIP-Seq data of Th1 cells. In order to identify differentiation-related chromatin changes, we also included our own H3K4me3 ChIP-Seq data for Thn cells. After we entered sample information into the system (**Supplementary Fig. 2a**), BioWardrobe downloaded the dataset and performed basic analysis (**Supplementary Fig. 2b**). ChIP-Seq data demonstrated the expected percentage of reads mapped and base frequency (**Supplementary Fig. 3a-d**), average tag density profiles showed high enrichment at promoters (**Supplementary Fig. 3ef**) and MACS2 identified large number of islands, the majority of which (68-77%) were located at promoters (**Supplementary Fig. 3g-j**). However, RNA-Seq results demonstrated poor mapping to the human transcriptome, poor coverage and potential DNA and ribosomal RNA contamination (**Supplementary Figs. 2** **and** **4, left side**). Keeping these problems in mind, we continued with data analysis and performed comparison of gene expression using DESeq2. Replicates were defined, genes were grouped by common transcription start site (TSS) and differentially expressed genes were identified. These results were used to define lists of genes that were expressed or silent in both Thn and Th1 cells or induced during differentiation. Next, H3K4me3 average tag density profiles were created for these three gene lists (**Fig. 1b**). As demonstrated in the graphs and Mann-Whitney-Wilcoxon (MWW) statistical analysis (**Fig. 1c**), genes that are expressed in both Thn and Th1 cells have higher levels of H3K4me3 at their promoters than genes that are silent in both cell types. Interestingly, differentiation-induced genes had intermediate levels of this modification in naïve cells, in which they were silent, suggesting that H3K4me3 poises inducible genes for expression during differentiation.

We used a second, published dataset to examine the role of KDM5B in regulating H3K4me in mouse embryonic stem (ES) cells (GSE53093 ^19^). The dataset includes RNA-Seq data for embryonic stem cells transfected with short hairpin RNA against luciferase (Control, shLuc) and against *Kdm5B* (shKdm5b) RNA. ChIP-Seq was performed for KDM5B and against H3K4 dimethylation and trimethylation (H3K4me2/3) in both shLuc- and shKdm5b ES cells. Basic analysis showed good quality datasets (**Supplementary Fig. 4, right** and not shown) for both ChIP-Seq and RNA-Seq. KDM5B was enriched at the promoters of expressed genes (**Fig. 2a**). H3K4me3 tag density profiles showed that *Kdm5b* knock-down resulted in a statistically significant redistribution of H3K4me3 tag density from promoters to the bodies of expressed genes (**Fig. 2b**). This was confirmed by heatmap analysis that showed spreading of both H3K4me2 (**Fig. 2c**) and me3 (not shown) into the gene bodies upon *Kdm5b* knockdown. Further, we compared H3K4me3 levels between individual H3K4me3 islands in shLuc- and shKDM5B-expressing cells using MAnorm. Interestingly, the majority of islands that have significantly (p < 0.01, fold > 2) gained H3K4me3 upon *Kdm5b* knock-down were located in gene bodies (**Fig. 2d**), confirming the results obtained from average tag density profiles and reported in the original publication.

**Figure 2.**
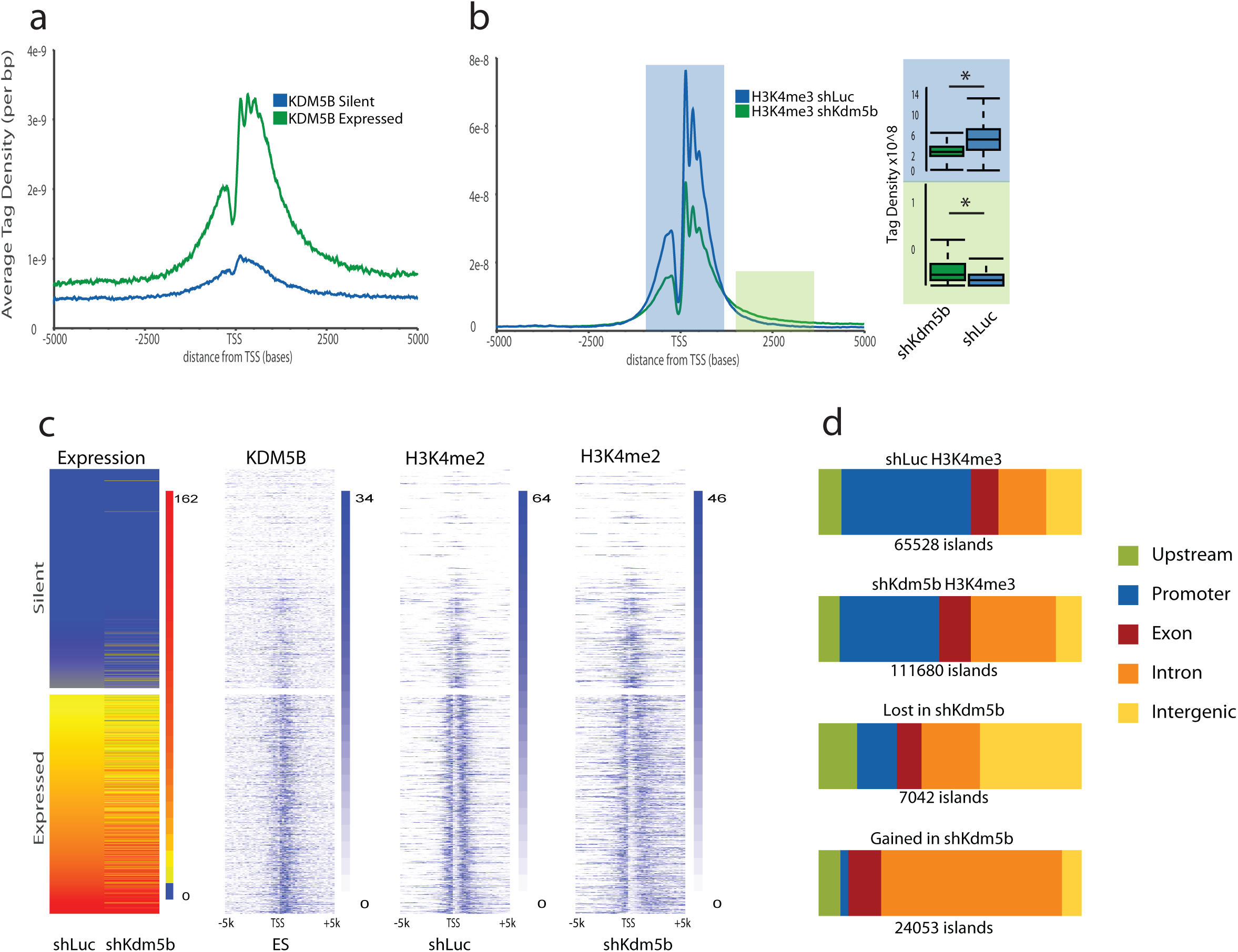
Role of KDM5B in the regulation of H3K4me3. **(a)** Tag density profile shows that KDM5B is recruited to promoters of expressed rather than silent genes in mouse ES cells. **(b)** *Kdm5b* knock-down results in the increase of H3K4me3 levels in the gene bodies and the corresponding loss of H3K4me3 at the TSSs of expressed genes. Box plots windows for the shaded areas are shown. **(c)** Heatmaps show that *Kdm5b* knock-down causes spreading of H3K4me2 into the gene bodies, but not upstream, of the expressed genes. **(d)** Genomic distribution of H3K4me3 peaks. Top graphs show data for all peaks in control (shLuc)- and shKdm5b-expressing cells; bottom graphs show distribution for those areas where occupancy was significantly (p < 0.01, fold > 4) increased or decreased upon *Kdm5b* knockdown.

In summary, we have developed a semi-automated system for storage, visualization and analysis of NGS data. BioWardrobe has been used to analyze data in several publications^20–24^. The system can be installed on Mac or Linux computers and can provide a data analysis solution for an entire laboratory or institution.

## Online Methods

### System overview

BioWardrobe allows users to upload, store and analyze NGS data. The workflow consists of two parts: basic and advanced analysis (**Supplementary Fig. 1**). The basic analysis includes operations that do not require comparison of samples: data download, quality control, calculation of RPKMs (reads per kilobase of transcript per million reads mapped), peak identification and upload to an integrated mirror of the UCSC genome browser. Advanced analysis includes comparing gene expression or chromatin immunoprecipitation sequencing (ChIP-Seq) profiles between samples. BioWardrobe can work with multiple genomes (our instance currently uses human, mouse rat, fly and frog) and additional genomes are easy to add, especially if the genome of interest is represented on UCSC genome browser. A flexible data ownership system is implemented: though all users can see all experiments on a local mirror of the UCSC genome browser, only members of the laboratories that own the data can access and analyze datasets within the BioWardrobe web interface or download it. Laboratory-level administrators can elect to share data with other laboratories. However, trusted bioinformaticians can have access to all datasets outside of BioWardrobe interface—e.g., via RStudio. We believe that this setup strikes a balance between maintaining data ownership and encouraging collaborations.

### Basic analysis

Basic analysis includes operations that are performed on a single library (**Supplementary Fig. 1a**). Analysis starts by entering the experiment description into Wardrobe. This information will be used to select the appropriate genome and analysis pipeline. Raw data can be directly downloaded by BioWardrobe via hypertext transfer protocol (http) or file transfer protocol (ftp) from core facilities or internet databases such as Gene Expression Omnibus (GEO) or Sequence Read Archive (SRA). Compressed or uncompressed FASTQ (.fastq) and SRA (.sra) files can be used. We elected not to use the prealigned BAM (.bam) files to ensure uniform alignment of samples.

For ChIP-Seq and similar experiments, reads are aligned to the genome with Bowtie ^12^, quality control analysis is conducted and data are summarized in a table (**Supplementary Fig. 2b**). In addition to basic statistics (percentages of mapped/unmapped/non-uniquely mapped reads and average fragment length), BioWardrobe displays several other quality control measures. Base frequency plots are used to estimate adapter contamination—a frequent occurrence in low-input ChIP-Seq experiments (**Supplementary Fig. 3c**). Average tag density profiles can be used to estimate ChIP enrichment for promoter proximal histone modifications (e.g., histone 3 lysine residue 4 trimethylation [H3K4me3], **Supplementary Fig. 3ef**). The genome browser can be used to visually compare results to other experiments in the database (**Supplementary Fig. 3gh**). ChIP-Seq results are displayed on the genome browser as coverage per million reads mapped. For paired-end reads, coverage is calculated as the number of fragments covering each base pair (bp). To obtain coverage for single-read experiments, average fragment length is calculated by model-based analysis of ChIP-Seq (MACS) ^15^, and individual reads are extended to this length in the 3’ direction. Islands (areas of enrichment) identified by MACS are displayed both on the browser (**Supplementary Fig. 3gh**) and as a table together with the nearest genes. The Island table can be used to select a cut-off for significant peaks that will be used in the downstream analysis. Additionally, the fasta sequences of peaks can be obtained with the click of a button and used with third-party tools (e.g., MEME-ChIP^25^) to produce sequence logos. Use of different parameters or pipelines for different antibodies (e.g., “broad peaks” MACS option for H3K27me3) is possible. Additionally, users can elect to use one of the experiments in the database as an “input” control for MACS. The distribution of the islands between genomic areas (promoters, exons, etc.) is displayed as a stacked bar graph (**Supplementary Fig. 3ik**).

For RNA sequencing (RNA-Seq) analysis, reads are aligned to the genome using RNA STAR (spliced transcripts alignment to a reference) ^13^ provided with the appropriate National Center for Biotechnology Information Reference Sequence (RefSeq) transcriptome. Other annotations can also be installed. The quality control tab displays the number of reads aligned within and outside the transcriptome. The percentage of the reads mappable to ribosomal DNA are displayed to estimate the quality of ribosomal RNA depletion (**Supplementary Fig. 2b**). Interpretation of quality control data is shown in **Supplementary Fig. 4**. Data are deposited on the browser, and RPKMs are calculated for each transcript (algorithm to be described elsewhere). Depending on the application, RPKM values can be presented for each transcript or summed up for each transcription start site (TSS) (for gene expression studies) or for each gene (for functional studies, e.g., Gene Ontology).

### Advanced analysis

If satisfied with the quality of data obtained from sequencing, a user can proceed to advanced analysis, which involves integration of information from multiple experiments. For gene expression analysis, the typical task is identifying differentially expressed genes. We elected to incorporate the DESeq1/2 algorithm ^16,26^ for this purpose because it does not require recreating transcript models and does not make many assumptions. In order to perform gene expression profiling, a user can define replicates and utilize the DESeq algorithm to calculate p-values and fold changes for all genes. On the basis of DESeq results, lists of genes whose expression changes can be created within BioWardrobe using expression levels, fold change, or p/q-values, as well as other parameters, and downloaded, if needed, in a table form for further analysis (e.g., gene set enrichment analysis).

The gene sets can also be used to create average tag density profiles and heatmaps within BioWardrobe (**Fig. 1b**). Average tag density profiles are used to compare the enrichment of histone modifications or other proteins around the TSS or the gene bodies between different gene sets. Often gene bodies used to estimate enrichment, for instance when comparing the levels of positive marks, such as H3K4me3, between expressed and silent genes. Heatmaps provide similar information but allow comparisons of modifications between individual genes. Statistical comparison of tag densities between groups of genes using Mann-Whitney-Wilcoxon (MWW) test can be performed by highlighting the area of interest with a mouse (**Fig. 1b, insert**). All graphs can be downloaded in publication-quality scalable vector graphics (SVG) format.

For ChIP-Seq, the task is usually the identification of areas that have different levels of binding between samples. The difficulty here is that the signal-to-background ratio (enrichment) is usually slightly different between ChIP-Seq experiments; thus, several assumptions have to be made in order to compare islands of enrichment. BioWardrobe uses the MAnorm algorithm ^17^, which assumes that modifications do not change in the majority of areas. This allows MAnorm to adjust for differential levels of enrichment between experiments. The lists of islands, fold changes and accompanying p-values are presented in table form, and islands can be viewed in the browser with the push of a button.

### R interface

Although, we sincerely believe that the set of quality control measurements and tools that we provide is the most useful, this may be a matter of personal preference. In order to allow for easy addition of custom analysis, we have incorporated R language script editing into BioWardrobe web interface for both basic and advanced analysis steps. System administrators can add custom R scripts in the R tab, and biologists can run these scripts via graphical web interface. In the basic analysis, customized R scripts can be run for each sample automatically or for selected samples. As an example, we have added scripts that provide the histogram of read pile-up or island length for ChIP-Seq data or gene body coverage and RPKM histogram for RNA-Seq data (**Supplementary Fig. 5**). In the advanced analysis R interface, customized scripts can be provided by system administrators. Users can select records of interest via the graphical user interface and run the customized scripts as needed. As an example, we provide a principal component analysis (PCA) script that can be used for analysis of RNA-Seq data (see http://demo.biowardrobe.com for output).

### Implementation

BioWardrobe is accessible via Google Chrome, Safari and Firefox browsers. The user interface is web based and utilizes HTML5 and JavaScript technologies. To speed up the development process, EXTJS and D3 JavaScript frameworks were used. On the server, Apache with PHP is used to process user’s requests. Linux or MacOSX native job schedulers are used to run Python pipelines. For stability, all pipelines have separate queues and process statuses. Pipeline output is stored in the SQL database with the exception of BAM files. These precomputed data are accessible by third-party software, like RStudio, that allows analysis that is not included in BioWardrobe. There are no specific hardware limitations for BioWardrobe. We have installed it on both a Linux server and Mac Pro desktop and laptop computers. An average Intel Core i7 computer with 32 gigabytes of RAM and a SATA HDD (more than 100 Mb read/write speed preferred) will analyze a typical ChIP-Seq or RNA-Seq experiment within less than 2 hours.

Current code and setup instructions are available at http://biowardrobe.com. A limited-functionality demo version is available at http://demo.biowardrobe.com.

## List of abbreviations used

ATAC-Seq: assay for transposase-accessible chromatin sequencing
ChIP: chromatin immunoprecipitation
ChIP-Seq: chromatin immunoprecipitation sequencing
D3: data-driven documents
DNase-Seq: DNase I hypersensitive sites sequencing
EXTJS: extension JavaScript
ftp: file transfer protocol
GEO: Gene Expression Omnibus
GRO-Seq: global run-on sequencing
HIPAA: Health Insurance Portability and Accountability Act of 1996
HTML: hypertext markup language
MACS: model-based analysis of ChIP-Seq
MAnorm: a model for quantitative comparison of ChIP-Seq datasets
MNase-Seq: micrococcal nuclease sequencing
MWW: Mann-Whitney-Wilcoxon
NGS: next-generation sequencing
PHP: PHP: hypertext preprocessor
RAM: random-access memory
RefSeq: National Center for Biotechnology Information Reference Sequence
RPKMs: reads per kilobase of transcript per million reads mapped
RNA-Seq: ribonucleic acid sequencing
SATA HDD: serial ATA hard disk drive
SRA: Sequence Read Archive (formerly known as Short Read Archive)
STAR: spliced transcripts alignment to a reference
SVG: scalable vector graphics
TSS: transcription start site(s)
Th1: T helper type 1
Thn: naïve T helper cells
UCSC: University of California, Santa Cruz

## Authors’ contributions

AK designed the system, AK wrote the program, AK and AB wrote the paper.

## Acknowledgements

Authors would like to thank Shawna Hottinger (Cincinnati Children’s Hospital Medical Center) for editorial assistance.

The work was supported in part by the Cincinnati Children’s Research Foundation and NHLBI NIH career transition award to A.B. (K22 HL098691).

---

**Supplementary Figure 1.**
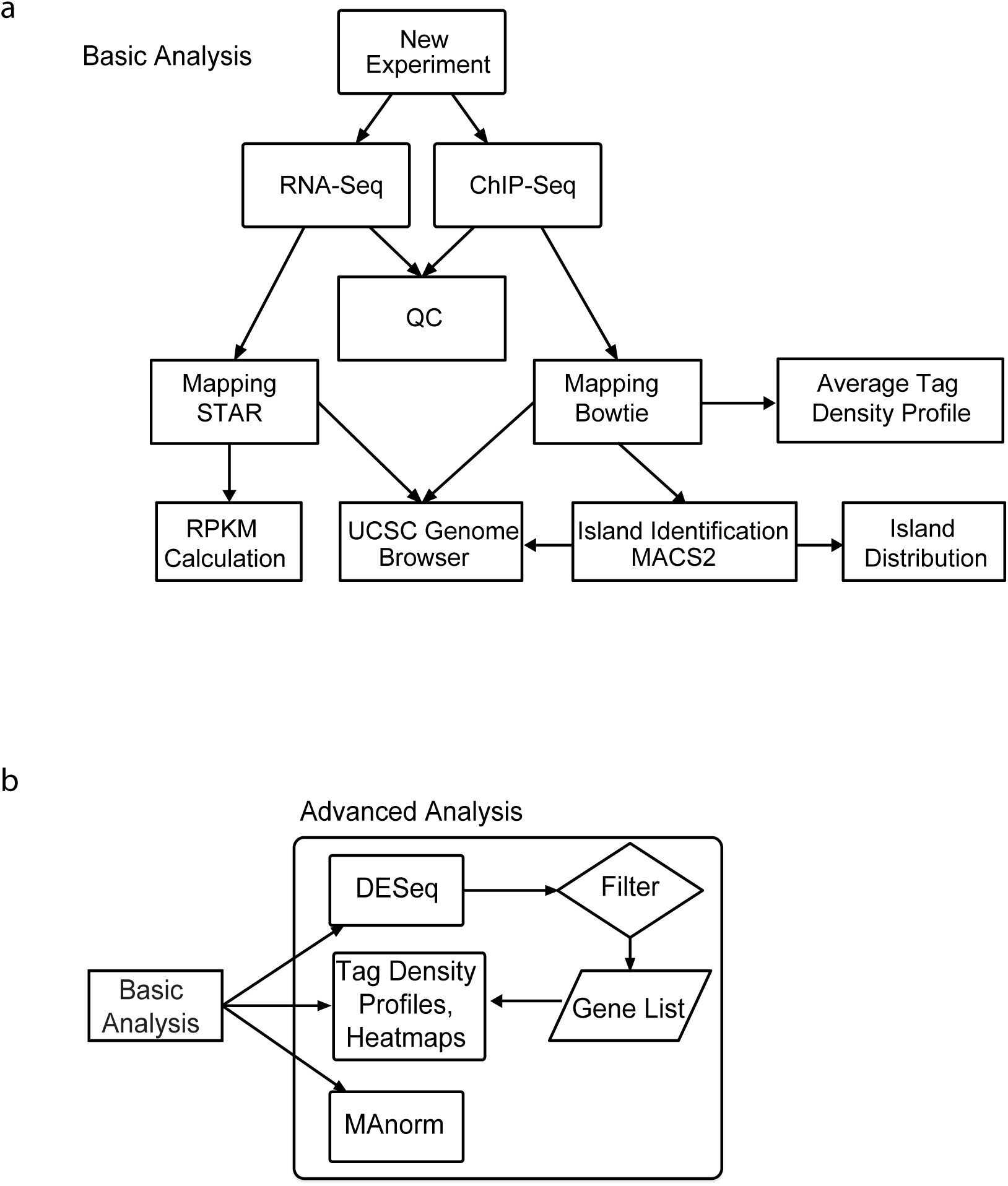
BioWardrobe pipelines. (**a**) Basic analysis pipelines. The flow diagram shows the tools used in the basic analysis pipelines for RNA-Seq and ChIP-Seq data. (**b**) Advanced analysis allows the user to identify differentially expressed genes using DESeq, create gene lists and use these lists to generate average tag density profiles and heatmaps. Differentially bound areas can be identified with MAnorm. QC, quality control.

**Supplementary Figure 2.**
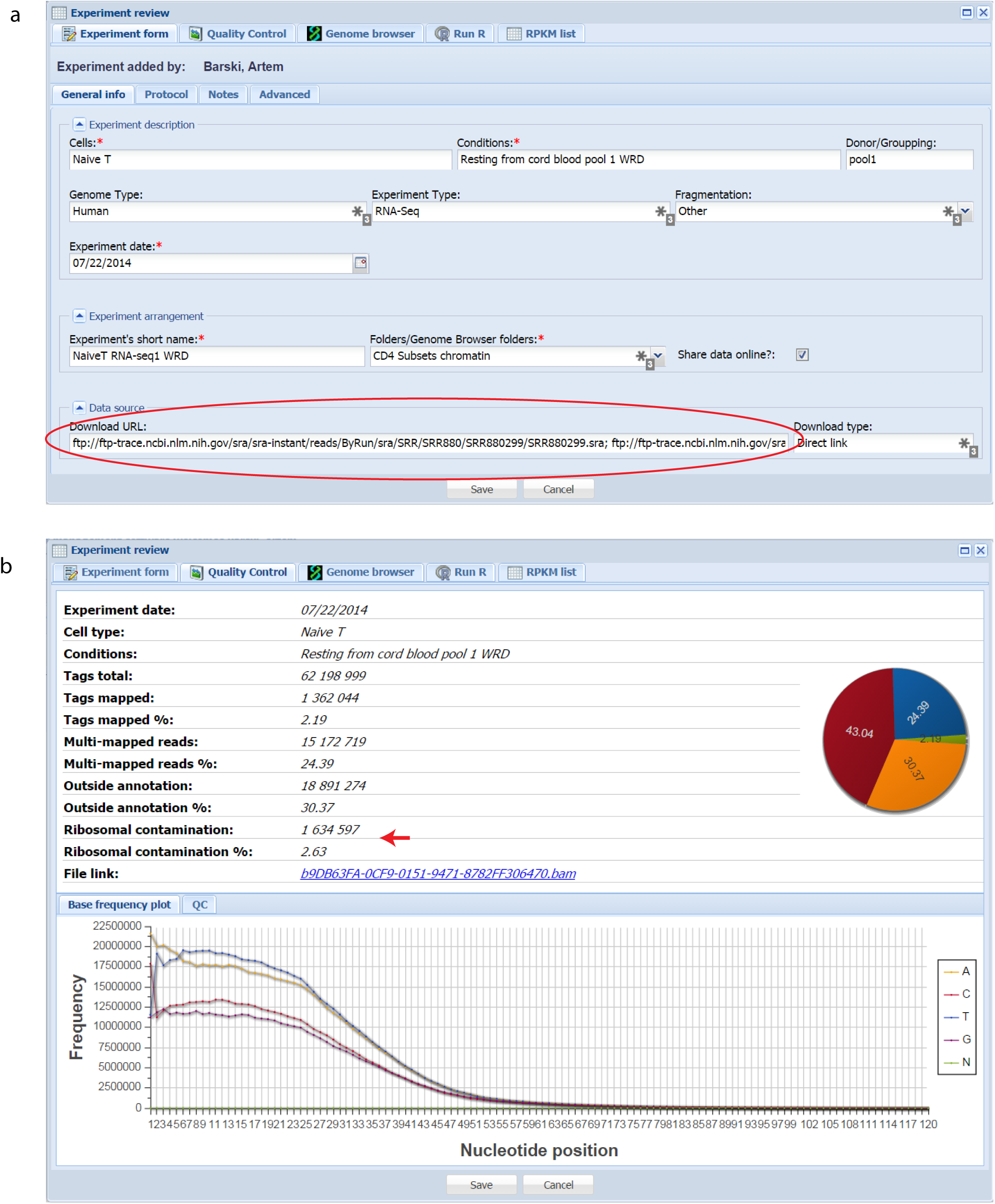
Experiment entry form and quality control window. (**a**) Experiments can be entered into Wardrobe by providing basic experimental details and a link to .fastq or .sra files. BioWardrobe will download the data, select the appropriate analysis pipeline and perform quality control. (**b**) Quality control tab shows basic mapping statistics. The arrow points to the high number of reads mappable to a ribosomal DNA repeat unit, suggesting incomplete removal of ribosomal RNA from the sample. This will not affect the results, but the experiment will require more sequencing since a large number of reads will be unproductively used on ribosomal RNA. Interpretation of other quality control measures is discussed in **Supplementary Figures 3** **and** **4**.

**Supplementary Figure 3.**
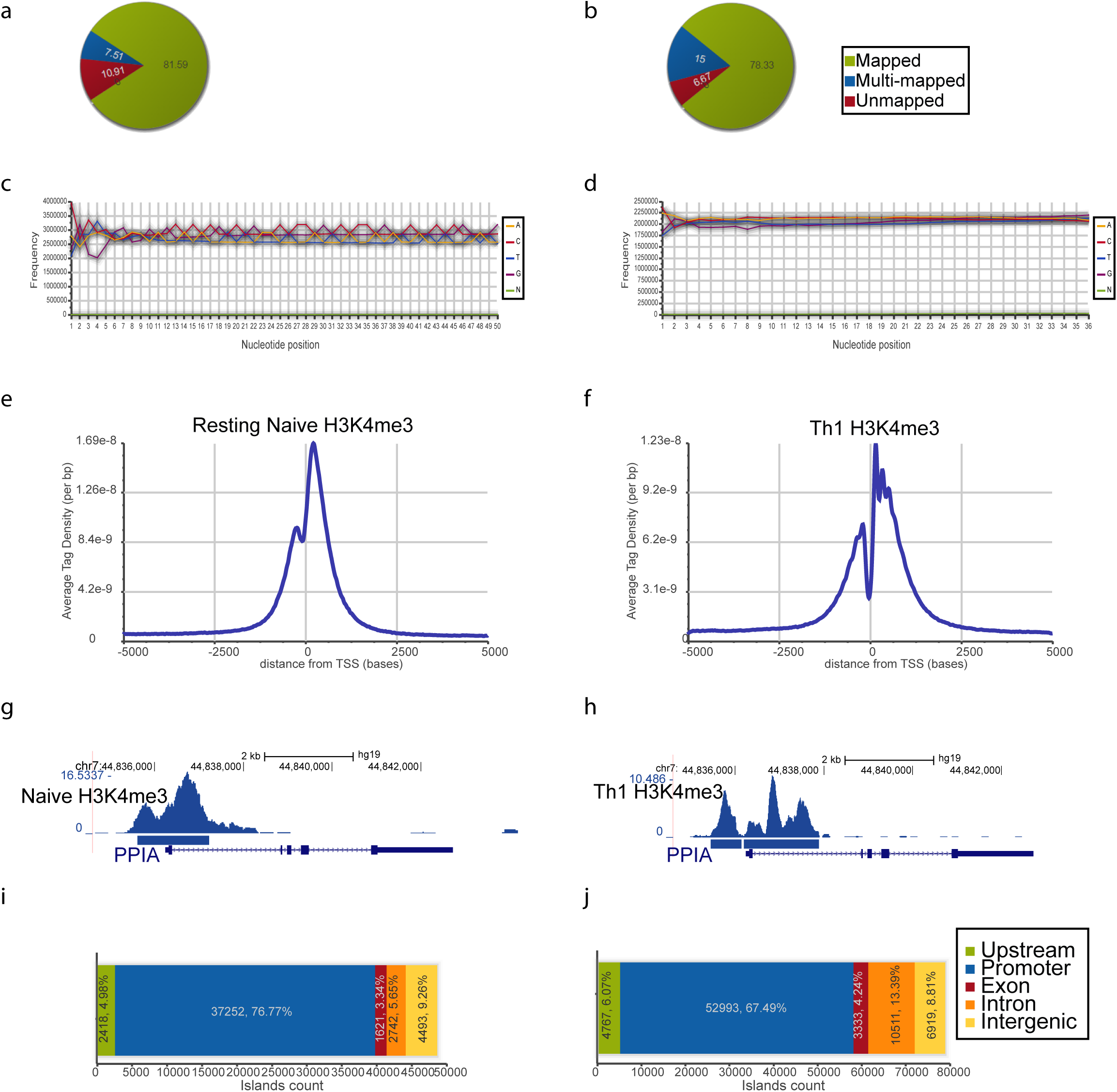
ChIP-Seq quality controls and interpretation. Data for H3K4me3 in Naïve (left) and Th1 cells (right) are displayed. (**a,b**) Pie charts show acceptable mapping statistics. (**c,d**) The absence of AT bias in these base frequency plots suggests enrichment of H3K4me3 in the vicinity of genic areas (also see **g**). The spiky plot in (**c**) is characteristic of adapter contamination in the library and suggests that the adapter/insert ratio during ligation needs to be decreased. This problem will not affect results, but the experiment will require more sequencing since a fraction of the reads will be unproductively used on adapter-dimers. (**e,f**) Average tag density profiles around the transcription start sites (TSSs) of all genes suggests that H3K4me3 is enriched around the TSS as expected. The experiment in (**e**) has slightly better enrichment, whereas the experiment in (**f**) has a better resolution due to a much shorter fragment size (estimated by MACS as 287 vs. 146 for **e** and **f**, respectively). (**g,h**) Representative browser images show H3K4me3 peaks at the *PPIA* promoter. Coverage by estimated fragments (top) and islands identified by MACS are shown. (**i,j**) BioWardrobe graphs show the distribution of H3K4me3 islands between genomic areas graphically and numerically (number of islands, percentage). Note that H3K4me3 is present primarily in promoter areas.

**Supplementary Figure 4.**
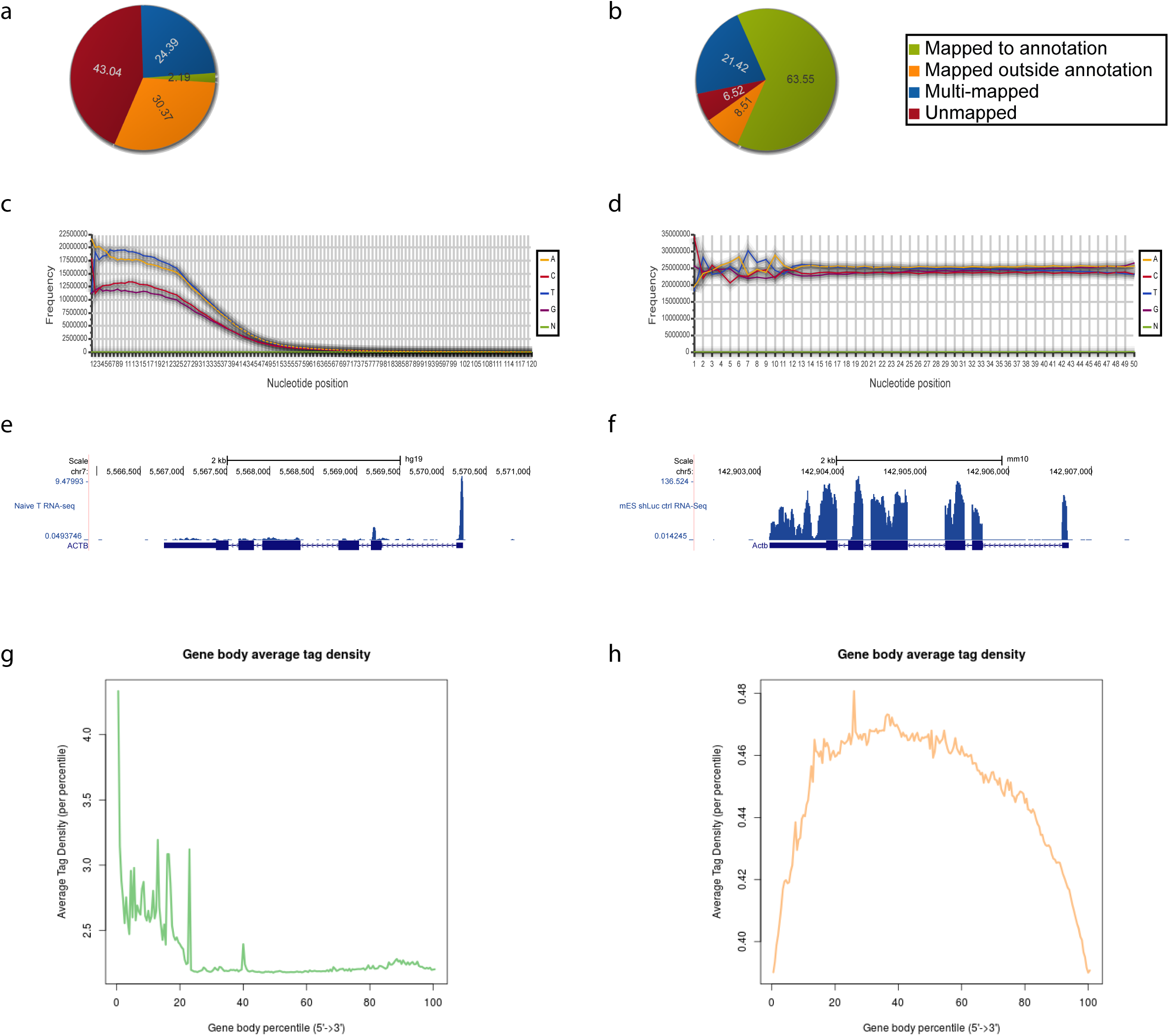
RNA-Seq quality controls and interpretation. RNA-seq data for Thn (left) and control mouse ES cells (right) are shown. (**a,b**) Pie charts show mapping statistics. In (**a**), note the poor mapping to transcriptome (high unmapped), large percentage of multi-mapped reads and reads mapping outside the annotation. These confirm insufficient ribosomal RNA removal (see also **Supplementary Figure 2b**) and suggest sample contamination with genomic DNA (or, less likely, a large amount of unannotated transcription). The latter is likely to inflate RPKM values for low expressed and non-expressed genes by a few units. (**c,d**) Base frequency plot. The general shape of the plot in (**c**) is characteristic of sequencing methods that produce reads of different lengths, such as Helicos. Also note a strong AT bias in (**c**). This bias is characteristic of genomic sequence but is not expected for transcripts, confirming the presence of DNA contamination in the sample. (**e,f**) Browser images show RNA-Seq coverage of *ACTB*/*Actb* genes. Note that the 5’ end of *ACTB* in (**e**) has a higher tag density (also see (**g**)). (**g,h**) Transcript coverage plots. Note that in (**g**) coverage is biased towards the 5’ end, suggesting either an inherent bias of the library construction method or RNA degradation.

**Supplementary Figure 5.**
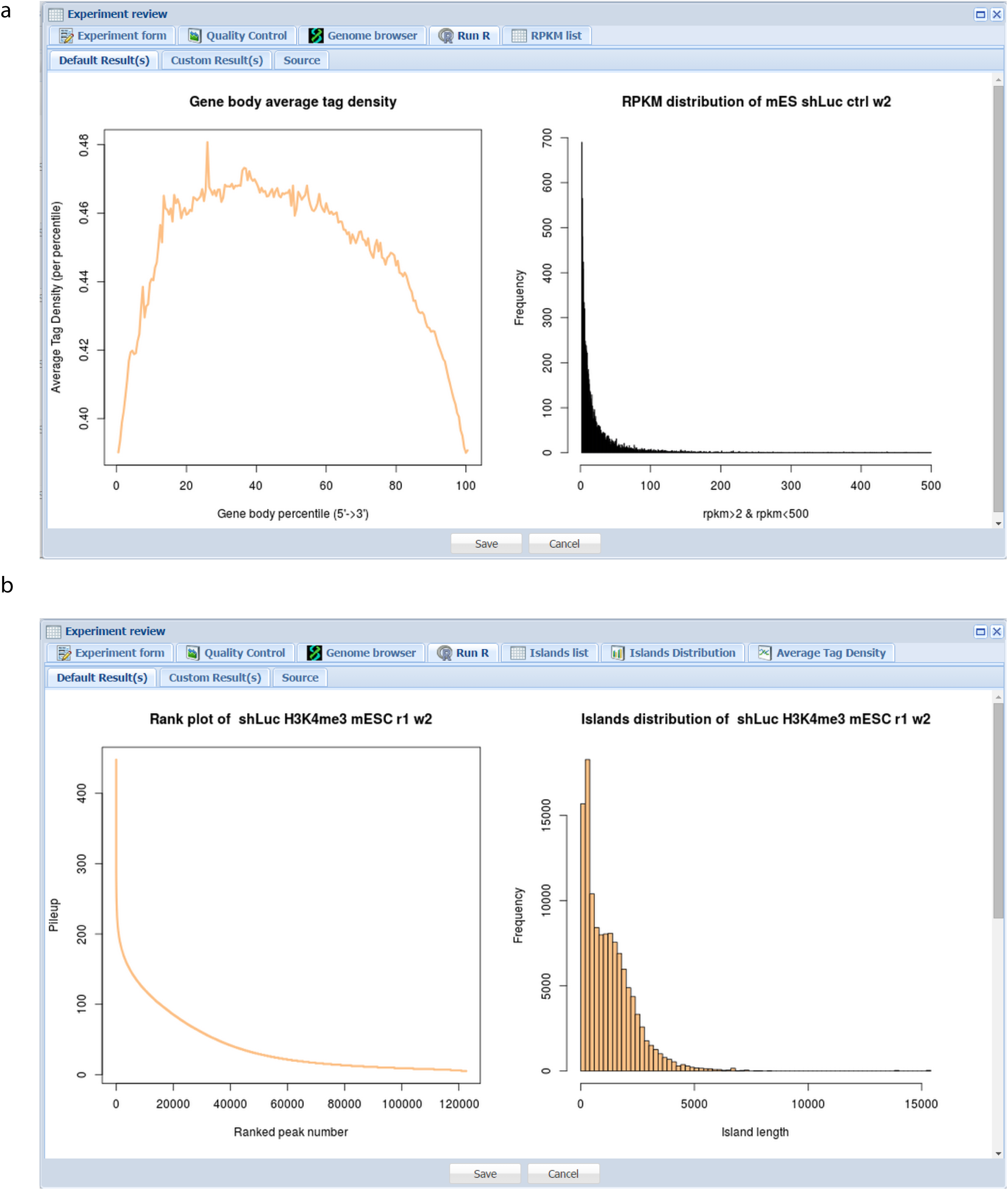
Example of plots produced by customizable R scripts. (**a**) Plots for RNA-Seq: gene body coverage and RPKM histogram. (**b**) Plots for ChIP-Seq: Rank vs. pile-up plot and Island length distribution plot.

